# Changes in antennal gene expression underlying sensory system maturation in *Rhodnius prolixus*

**DOI:** 10.1101/2021.09.02.458747

**Authors:** Jose Manuel Latorre-Estivalis, Ewald Große-Wilde, Gabriel da Rocha Fernandes, Bill S. Hansson, Marcelo Gustavo Lorenzo

**Affiliations:** Instituto de Fisiología, Biología Molecular y Neurociencias, Universidad de Buenos Aires - CONICET, Buenos Aires, Argentina; Department of Evolutionary Neuroethology, Max Planck Institute for Chemical Ecology, Jena, Germany; Faculty of Forestry & Wood Sci, Excellent Team for Mitigation, Czech University Life Sci, Prague, Czech Republic; Plataforma de Bioinformática, Instituto René Rachou - FIOCRUZ, Belo Horizonte, Minas Gerais, Brazil; Vector Behavior and Pathogen Interaction Group, Instituto René Rachou - FIOCRUZ, Belo Horizonte, Minas Gerais, Brazil

**Keywords:** Age, transcriptome, antennae, *Rhodnius prolixus*, sensory genes

## Abstract

**Background:** Triatomine bugs are the blood feeding insect vectors transmitting Chagas disease to humans, a neglected tropical disease that affects over 8 million people, mainly in Latin America. The behavioral responses to host cues and bug signals in *Rhodnius prolixus* are state dependent, i.e., they vary as a function of post-ecdysis age. At the molecular level, these changes in behavior are probably due to a modulation of peripheral and central processes. In the present study, we report a significant modulation of the expression of a large set of sensory-related genes. Results were generated by means of antennal transcriptomes of 5th instar larvae along the first week (days 0, 2, 4, 6 and 8) after ecdysis sequenced using the Illumina platform.

**Results:** Age induced significant changes in transcript abundance were established in more than 6,120 genes (54,7 % of 11,186 genes expressed) in the *R. prolixus* antenna. This was especially true between the first two days after ecdysis when more than 2,500 genes had their expression significantly altered. In contrast, expression profiles were almost identical between day 6 and 8, with only a few genes showing significant modulation of their expression. A total of 86 sensory receptors, odorant carriers and odorant degrading enzymes were significantly modulated across age points and clustered into three distinct expression profiles.

**Conclusions:** The set of sensory genes whose expression increased with age (profile 3) may include candidates underlying the increased responsiveness to host cues shown by *R. prolixus* during the first days after molting. For the first time, we describe the maturation process undergone at the molecular level by the peripheral sensory system is described in an hemimetabolous insect.

## Background

Sensory performance is a fundamental component underlying the behavioral repertoire of an organism. It defines how precisely an animal can keep track of relevant events happening in its environment. The sensory sounding of the *umwelt* of insects is predominantly multimodal, i.e., the world is accessed through diverse senses and integration of sensory inputs frequently drives behavior [1, 2]. This seems particularly true for model kissing-bugs like *Triatoma infestans* and *Rhodnius prolixus*, which vector *Trypanosoma cruzi*, causative agent of Chagas disease, to humans [3]. Sensory neurons (SNs) covering diverse modalities concentrate mainly in the antennae of insects. These include olfactory, gustatory, thermo-, hygro-, or mechanosensory units that are finely tuned to detect their cognate stimuli. Nevertheless, the acuity and sensitivity of these sensory systems is not a static feature. As an example, it is well known that antennal responsiveness to odors presents a circadian variation in insects [4–7].

Insect behavioral responsiveness to communication signals and host cues can depend on factors such as age, nutritional or reproductive status. The responsiveness of sensory components informing decision-making neuropiles in the central nervous system is also affected by these factors. As an example, the imaginal moult involves a dramatic transformation of the insect that normally requires a few hours/days of final adjustments after ecdysis. This process is usually considered as a maturation process affecting behavior. Indeed, recently emerged mosquito females do not engage in host seeking, a behavior only expressed between the second and third day after their imaginal moult [8, 9]. Insects that engorge on blood also commonly show dramatic changes in their behavior due to the physiological processes abruptly triggered by the blood meal [10, 11]. After biting a host and becoming fully-engorged, blood-sucking insects quit host-seeking almost immediately [12]. In the case of mosquito females, two days later they initiate the search of a breeding site where to lay the egg batch [13]. Considering that such dramatic changes affecting foraging activities can be established, it seems feasible that they have molecular correlates both at the sensory periphery and in central processing centers.

While the detection of cues related to meal sources or oviposition sites is transiently needed for nurture or egg-laying, the detection of many other clues and signals needs to be permanently effective. This is the case for the recognition of predator-emitted odors, aversive compounds present in non-edible plants, alarm and aggregation pheromones. Some triatomines use aggregation signals present in their feces to mark their hiding places [14, 15]. Aggregation responses to these signals are seen in all instars independently of age [16] and under most nutritional conditions [17]. Another example are the alarm signals emitted by adult triatomines when exposed to mechanical disturbance [18], as all instars need to present avoidance reactions to these pheromones in order to stay away from danger independently of their physiological condition. Therefore, one would expect that the molecular machinery committed to detect this type of stimuli should not vary significantly with age, neither should it change with nutritional status.

The developmental stage, age, and nutritional, and reproductive status of insects can affect SN sensitivity (reviewed by [19]). This is quite clear for kissing-bugs, insects that feed very sporadically on the blood of vertebrates that they steal in the still of the night. As an example, the antennae of *R. prolixus* are known to respond to ammonia with increasing electroantennogram amplitudes depending on starvation, but this is only true during the scotophase [20]. Adult mosquitoes also present significant changes in SN responsiveness as a function of post-emergence age [8, 21–23]. Peaks of responsiveness by SNs may also depend on reproductive status as observed in moths [24, 25]. Finally, it was recently shown that Dengue I infection can induce augmented olfactory responses in the antennae of infective mosquitoes [26], suggesting that pathogens can also modulate OSNs responsiveness.

Reports on gene expression profiles as a function of age, nutritional or mating status have proven that insect antennae go through a complex set of processes regulating the transcript abundance of diverse genes. Mosquito age after adult emergence affects the transcriptomic profile seen in the antennae of *Ae. aegypti*, with many sensory receptor-coding genes showing changes in their expression over the first five days of adult life in both sexes [8]. Furthermore, similar regulation of gene expression profiles in the antennae of females of the species is affected through the temporal interval of the first gonotrophic cycle and after ingestion of a blood meal [27]. Females of the malaria vector *Anopheles gambiae* also show clearly regulated expression changes in their antennae as a consequence of engorging on blood [28]. In the antennae of the hemimetabolous vector bug *R. prolixus*, the imaginal moult triggers a clear increase in the expression of many chemosensory genes compared to the last larval instar [29] suggesting that the antennae of adults become more sensitive to diverse odor molecules.

As Bodin et al. [30] have shown, triatomine responsiveness to host cues depends on age after ecdysis. While recently molted bugs are not attracted towards host-mimicking heat sources or CO_2_-added air currents, they show consistent responses to these host-related cues a week later. To date, it is not clear whether this is an exclusive consequence of a maturation process affecting peripheral and/or central components controlling responsiveness to host cues. We hypothesize that the abundance of olfactory, gustatory, thermo- and hygroreceptors potentially mediating the detection of host-related cues may vary through the maturation interval. In addition, we suggest that receptor proteins dedicated to the detection of aggregation and alarm pheromones should present a stable level of expression through the same interval. Our work aims to elucidate the extent to which sensory gene expression profiles change in 5^th^ instar bug antennae as a proxy to understand kissing-bug sensory plasticity at the peripheral level through the post-ecdysis maturation of bug behavior.

## Results

### Exploratory analysis

Samples from the five age points presented different expression profiles according to the PERMANOVA analysis plot (Fig. 1), indicating a strong effect of age on the sample distribution (PERMANOVA, R^2^ =0.42, corrected p-value <0.01, Additional file 1: Table S1). In Fig. 1, replicates from days 0 and 2 appeared completely separated and paired comparisons with the other time points were all significant (Additional file 1: Table S1). On the other hand, replicates from days 4, 6 and 8 grouped together, with only days 4 and 8 differing significantly (Fig. 1 and Additional file 1: Table S1).

**Fig. 1.**
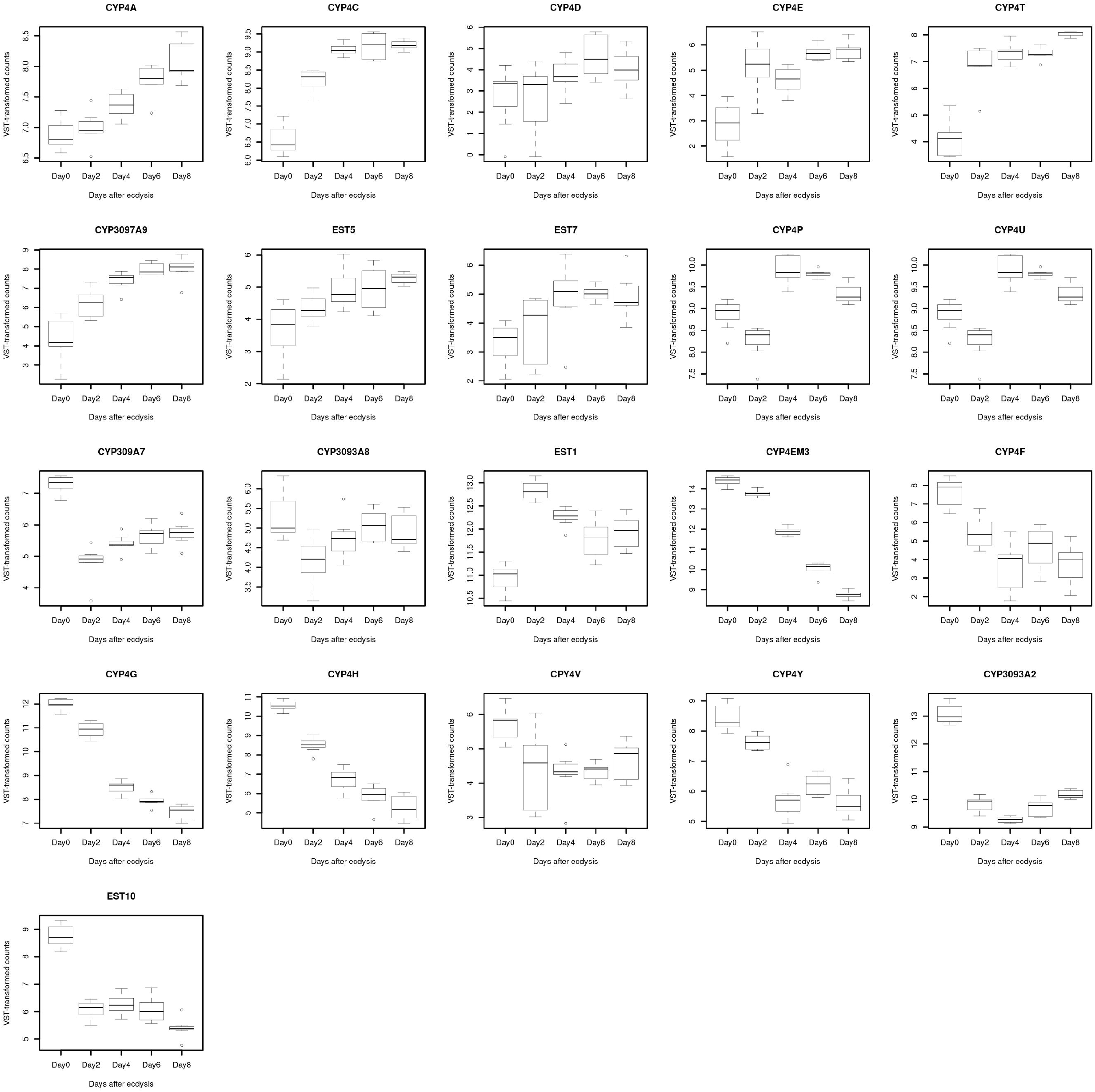
Principal Coordinates Analysis Plot. The plot was created using the *ade4* R package. The impact of the variable “age” on the distances was measured by a PERMANOVA analysis using 999 permutations in R.

### Overall expression changes

According to the GLM analysis performed in EdgeR, a total of 6,120 transcripts changed their antennal expression significantly during the first week of *R. prolixus* larval life. This represents 54,7 % of 11,186 transcripts (after the HTSFilter processing step). The differentially expressed genes (DEGs) were identified based on False Discovery Rate (FDR) <0.05 with no fold-change threshold (Additional file 2: Table S2).

### Highly expressed sensory genes along age

A consensus of 43 target genes was obtained from the top 50 most expressed sensory genes in each time point (Additional file 3: Table S3). As expected, odorant binding proteins (OBPs) and chemosensory proteins (CSPs) were the most abundant within this set, with 18 and 14 members, respectively. Regarding the other protein families, few members were highly expressed: *RproGr27, RproIr93, RproSnmp1a, RproPain, RproAmmT1*, 4 odorant degrading enzymes (ODEs) and the PPK *Rprrc013510*.

### Pairwise comparisons

Expression data from the different time points was compared using a pairwise analysis to identify ages at which the transcriptional changes occurred (Table 1). A total of 3,290 genes changed their expression significantly (*s*-values <0.05 and on an absolute fold-change threshold >1.5) in at least one of the comparisons performed. As we observed in the MA and PCoA plots, the greatest change in antennal expression occurred right after day 0 (with a total of 2,534 genes showing significant changes compared to a pool of the remaining days). The second greatest gene expression change was detected between day 0 *vs*. day 2 (1,738 genes representing 16,27%). These transcriptional changes are reflected in the corresponding MA-plots (Additional files 4 and 5: Figs. S1 and S2). The numbers of DEGs in the antennae decreased between subsequent ages, as well as the amplitude of their changes (Table 1). This reduction was more pronounced between days 6 and 8, when only 80 genes changed their expression within a smaller range, supporting the clustering pattern observed in the MA and PCoA plots (Fig. 1).

**Table 1.**
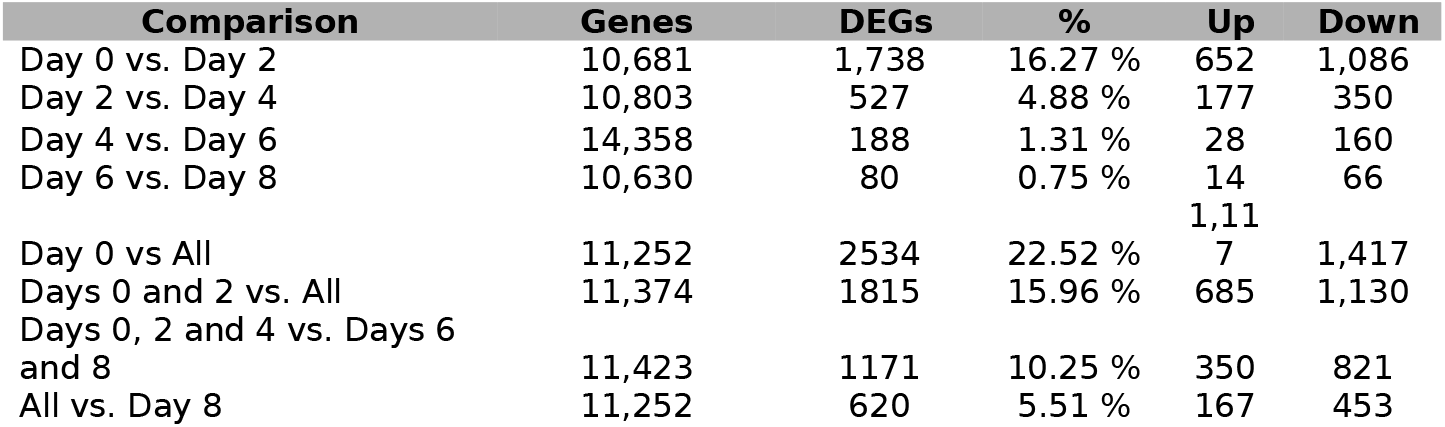
Number of differentially expressed genes identified in different time point comparisons performed along eight days after ecdysis. DEGs: Differentially expressed genes.

When the direction of the transcriptional changes observed in the antennae was analyzed, we observed that the number of down-regulated genes was higher than that of up-regulated ones in all comparisons (Table 1). This was highest at older ages when 82% of DEGs decreased their expression, while it was lower than 66% during the first four days. All MA-plots reflected how down-regulation is predominant over upregulation in all comparisons (Additional files 4 and 5: Figs. S1 and S2).

A total of 86 sensory receptors, odorant carriers (OBPs and CSPs) and ODEs were identified as differentially expressed (Table 1 and Additional file 6: Table S4). Eighteen out of 116 OR genes had their expression significantly altered, representing 15% of the OR set. Interestingly, more than 30% of all transient receptor potential (TRP) channels presented significant changes in their expression profiles (5 out of 14) and a similar case was observed for pickpockets (PPKs) (3 out of 10). In contrast, only two IRs and two GRs were differentially expressed, i.e., less than 7% of each family showed modulated expression. The expression of most CSPs was affected by age, 15 out of 18 were found to be differentially expressed. For genes belonging to the OBP and ODE families, the percentage of DEGs was also high, with 13 (46%) and 21 (65%) members changing their expression during the first week after ecdysis.

The temporal pattern of gene expression observed was consistent with the overall changes described above, with the greatest alteration occurring between day 0 *vs*. day 2 (49 DEGs) followed by a decrease in the number of DEGs in subsequent comparisons (15, 7 and 3 DEGs, respectively). This situation was extremely clear with sensory receptors that varied almost exclusively between the first two days (16 DEGs). After day 2 only one odorant receptor (OR) and several sensory neuron membrane proteins (SNMPs) displayed changes in their expression levels (Table 2). In the case of odorant carriers and ODEs, the decrease observed in the number of DEGs progressed across all time points (Table 2). A higher number of targets were identified as DEGs in the grouped comparisons, suggesting that changes in some genes are cumulative and could not be detected by sequential comparisons. Down-regulation was always more frequent in all comparisons, except for the one between days 2 and 6 when up-regulated genes were more abundant (Table 2 and Additional file 6: Table S4).

**Table 2.** Number of differentially expressed genes from the sensory gene families identified in different temporal comparisons performed along eight days post-ecdysis. DEGs: differentially expressed genes; ORs: odorant receptors; GRs: gustatory receptors; IRs: ionotropic receptors; TRPs: transient receptor potential channels; SNMPs: sensory neuron membrane proteins; CHEB: Chemosensory proteins B; AmmT: Ammonium transporter; CSPs: Chemosensory proteins; OBPs: odorant binding proteins; ODEs: Odorant degrading enzymes. ODEs include cytochromes and esterases.

| Gene Family (DEGs) | Day 0<br>vs.<br>Day 2 | Day 2<br>vs.<br>Day 4 | Day 4<br>vs.<br>Day 6 | Day 6<br>vs.<br>Day 8 | Day 0<br>vs.<br>All | Days 0-2<br>vs.<br>All | Days 0, 2, 4<br>vs.<br>Days 6 and<br>8 | All<br>vs.<br>Day<br>8 |
| --- | --- | --- | --- | --- | --- | --- | --- | --- |
| ORs (18) | 7 | 0 | 0 | 1 | 16 | 4 | 2 | 2 |
| GRs (2) | 0 | 0 | 0 | 0 | 1 | 0 | 1 | 0 |
| IRs (2) | 2 | 0 | 0 | 0 | 2 | 2 | 0 | 0 |
| TRPs (5) | 4 | 0 | 0 | 0 | 4 | 3 | 2 | 0 |
| PPKs (3) | 2 | 0 | 0 | 0 | 1 | 1 | 0 | 0 |
| SNMPs (3) | 1 | 1 | 1 | 0 | 1 | 1 | 1 | 2 |
| CHEB (1) | 0 | 0 | 0 | 0 | 1 | 0 | 0 | 0 |
| AmmT (1) | 0 | 0 | 0 | 0 | 1 | 0 | 0 | 0 |
| CSPs (15) | 10 | 4 | 3 | 1 | 11 | 10 | 9 | 6 |
| OBPs (13) | 9 | 2 | 1 | 0 | 9 | 7 | 6 | 5 |
| ODEs (21) | 14 | 8 | 2 | 1 | 17 | 17 | 13 | 7 |
| **Total (84)** | **49** | **15** | **7** | **3** | **64** | **45** | **34** | **22** |
| Receptors (35) | 16 | 1 | 1 | 1 | 27 | 11 | 6 | 4 |
| Carriers and ODEs (49) | 33 | 14 | 6 | 2 | 37 | 34 | 28 | 18 |
| Up | 14 | 9 | 5 | 0 | 30 | 18 | 11 | 6 |
| Down | 35 | 6 | 2 | 3 | 34 | 27 | 23 | 16 |

### Expression changes of odorant, ionotropic and gustatory receptors: examples of chemosensation regulated at the periphery

A total of 18 ORs changed their antennal expression significantly during the first week after ecdysis (Additional file 6: Table S4). These receptors clustered in three profiles according to their expression (Fig. 1). The first (profile 1) was composed of four genes (*RproOr17, RproOr39, RproOr96*, and *RproOr104*), whose expression decreased with age. Fig. 3 depicts this clearly for *RproOr17* and *RproOr96*, but also for *RproOr39* and *RproOr104*, whose expression decreased significantly on day 2. The second cluster defined a smaller group of genes that showed a significant down-regulation mostly after day 6 (profile 2). *RproOr100* was the only OR in the second cluster (Fig. 2) and presented a significantly decreased expression on day 8 (Fig. 3). The last profile is composed of 13 ORs (Fig. 2), that increased their expression with bug age (profile 3). These genes presented two distinct patterns (Fig. 3): either a continuous rise in expression through the whole interval, e.g., *RproOr30* and *RproOr72*, or an augmented expression at day 2 followed by stabilization, e.g., *RproOr7* and *RproOr18*.

**Fig. 2.**
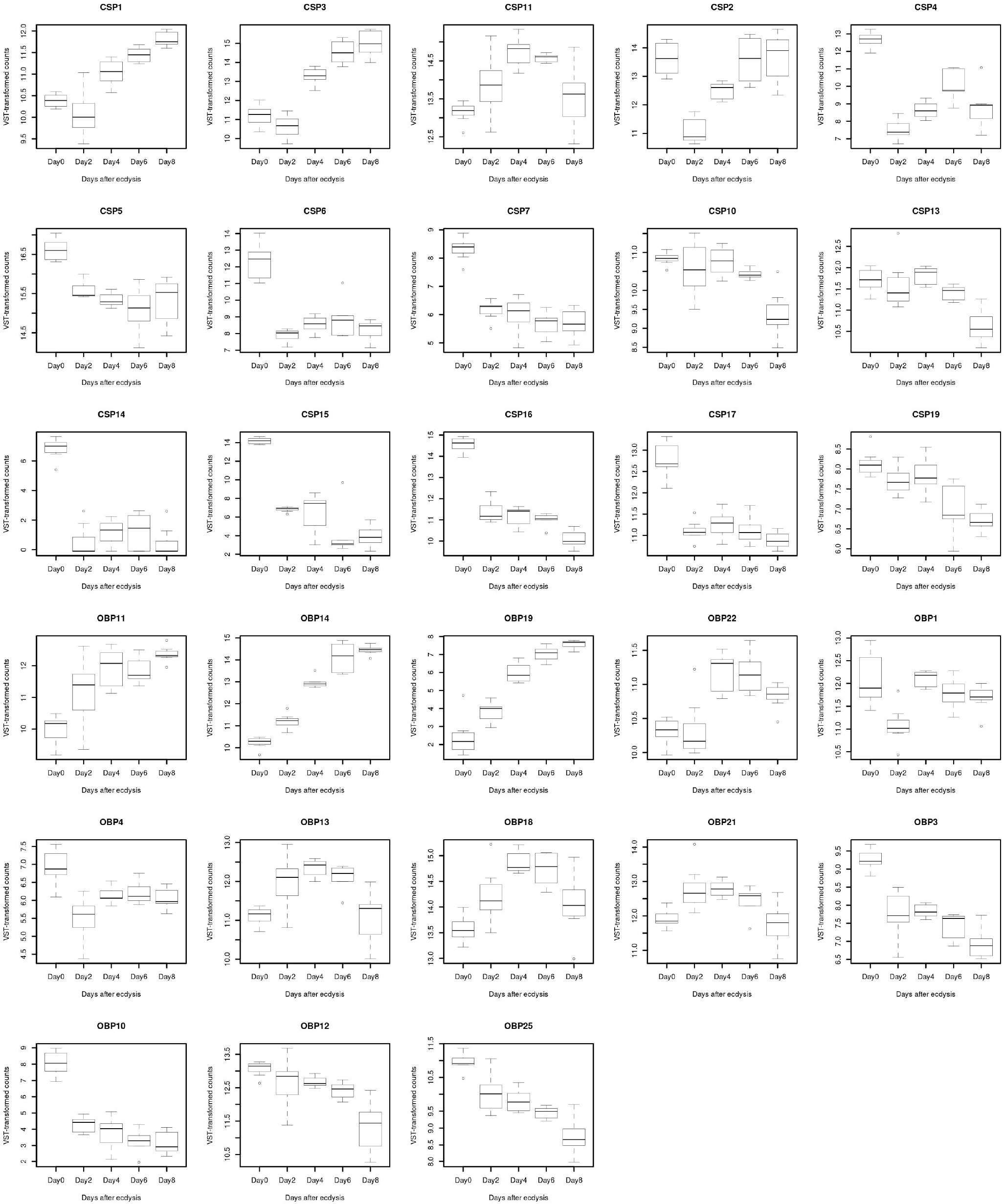
Effect of age on the antennal transcription of sensory-related genes. The heatmap was created using transformed count data (generated with *vst* function in DESeq2 and included in Additional file 10: Table S8) as input of the pheatmap R package. This calculated a z-score for each gene and plotted it by means of a color scale, where blue/red represent lowest/highest expression. Dendrogram was created with hierarchical clustering among genes based on their Euclidean distances and the Ward method for clustering.

**Fig. 3.**
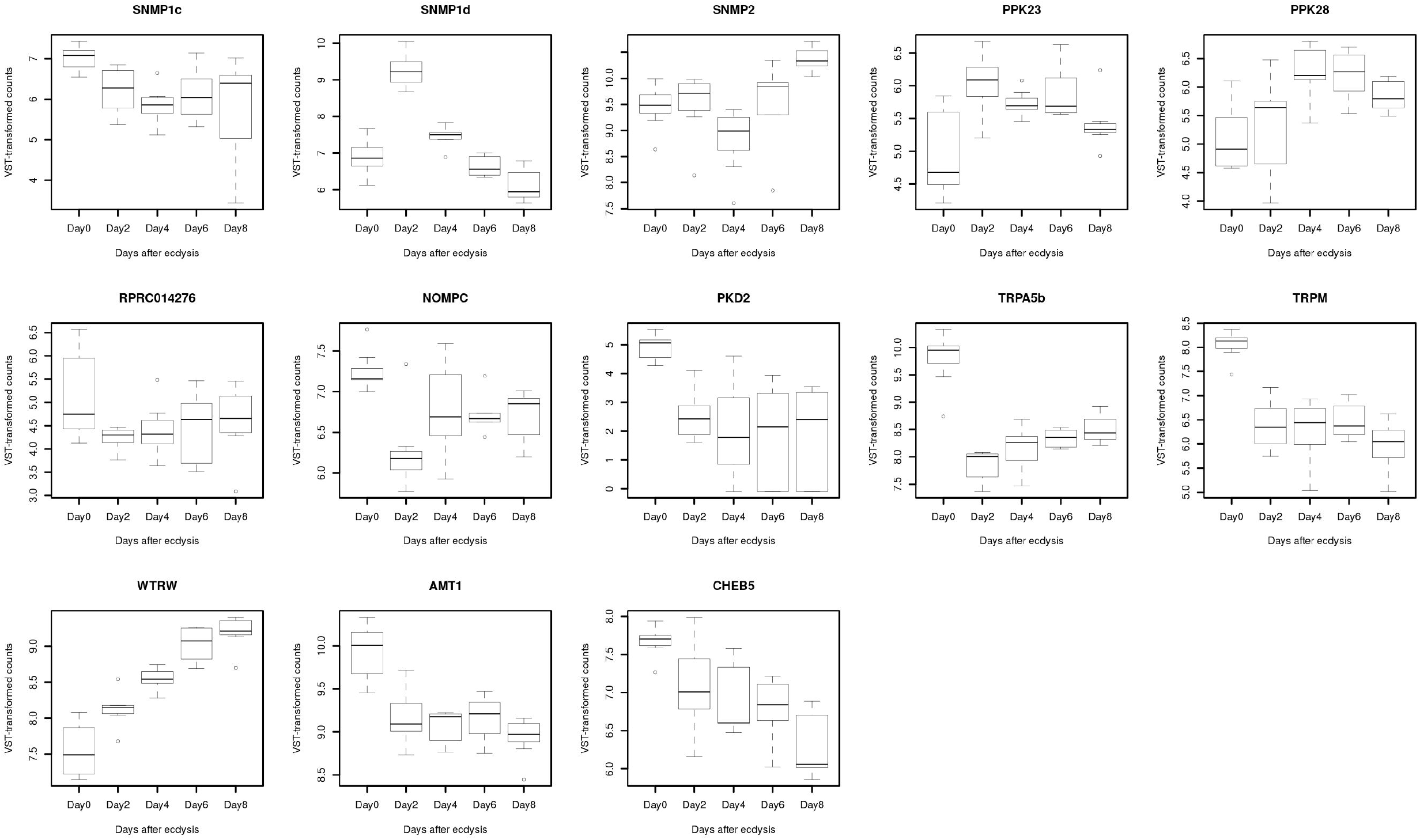
Box-plots of differentially expressed odorant, ionotropic and gustatory receptors along five age points post-ecdysis. Gene expression is represented as Variance Stabilizing Transformed (VST) counts that were obtained from count data and the *vst* function from DESeq2. VST counts are reported in Additional file 10: Table S8.

Transcriptional changes were very restricted for IRs and GRs, as only four of these genes changed their expression significantly (Fig. 3 and Additional file 6: Table S4). Two ionotropic receptors (IRs) belonging to profile 1, i.e. *RproIr93a* and *RproIr107*, decreased their expression drastically between days 0 and 2 (Fig. 3). A similar case was seen for the gustatory receptor (GR) *RproGr14* that showed a significant decrease after day 4 (Fig. 3). Interestingly, the expression of *RproGr26* rose steadily until day 6 (profile 3).

### Expression changes of PPK and TRP receptors

The expression of only three PPKs receptors was affected by age (Fig. 2 and Additional file 6: Table S4). *Rproppk23* and *Rproppk28* belonging to profile 1 showed increased expression at the first age points (Fig. 4a). The expression pattern of the other PPK (*Rprc014276*) fit to profile 1, showing a reduction on day 2 (Fig. 4a).

**Fig 4.**
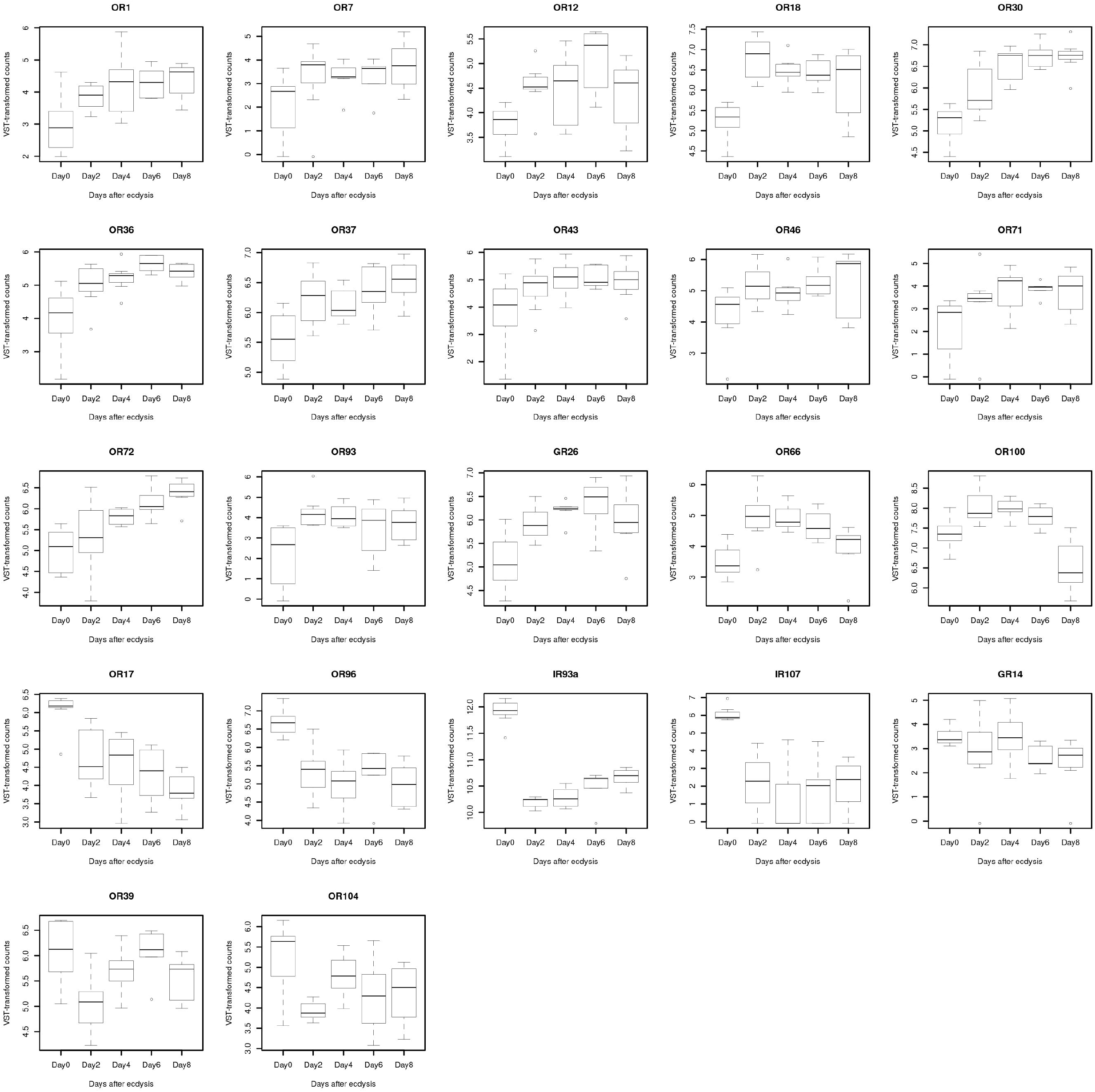
Box-plots of differentially expressed pickpocket, transient potential receptor channels and other sensory receptors along five age points postecdysis. Gene expression is represented as Variance Stabilizing Transformed (VST) counts that were obtained from count data and the *vst* function from DESeq2. VST counts are reported in Additional file 10: Table S8.

Two different groups of TRP channels were identified by the clustering analysis according to their expression profiles (Fig. 2). *RproNompC*, *RproPKD2*, *RproTrpA5b* and *RproTrpM* belonged to profile 1, exhibiting a significant reduction on day 2 (Fig. 4b and Additional file 6: Table S4). In contrast, *Rprowtrw* showed increased expression through the first half of the age interval, being clearly associated with profile 3 (Fig. 4b and Additional file 6: Table S4).

### Other membrane sensory proteins

Three SNMPs were identified as DEGs distributed in all three clusters (Fig. 2 and Additional file 6: Table S4): *RproSnmp1c* (profile 1), which showed a decline until day 4; *RproSnmp1d* (profile 2), with a remarkable peak of expression at day 2 followed by a pronounced reduction through the following days; and *RproSnmp2* (profile 3) with a significant increase after day 6. Two additional genes clustered in profile 1: ammonium transporter 1 (*RproAmt1*) and the chemosensory protein B5 (*RproCheB5*), which presented a significantly decreased expression after day 0 (Fig. 4c).

### Expression changes of odorant carriers

The effect of age on CSPs and OBPs was very clear, with 83% and 48% of them being differentially expressed, respectively (Additional file 6: Table S4). A first group including 10 CSPs and 5 OBPs that fit into profile 1 had its expression reduced with age (Fig. 2). Dramatic examples of reduced expression could be seen with *RproCsp6*, *RproCsp14, RproCsp17 or RproObp25* (Fig. 5). Three genes fit into profile 2, i.e., *RproObp12, RproCsp10* and *RproCsp13*, showing a significant reduction only after day 6 (Fig. 5). Profile 3 was seen for 3 CSPs and 7 OBPs (Fig. 2) that displayed a rise in expression with age, as seen for *RproCsp1, RproObp11, RproObp14* and *RproObp19* (Fig. 5).

**Fig. 5.**
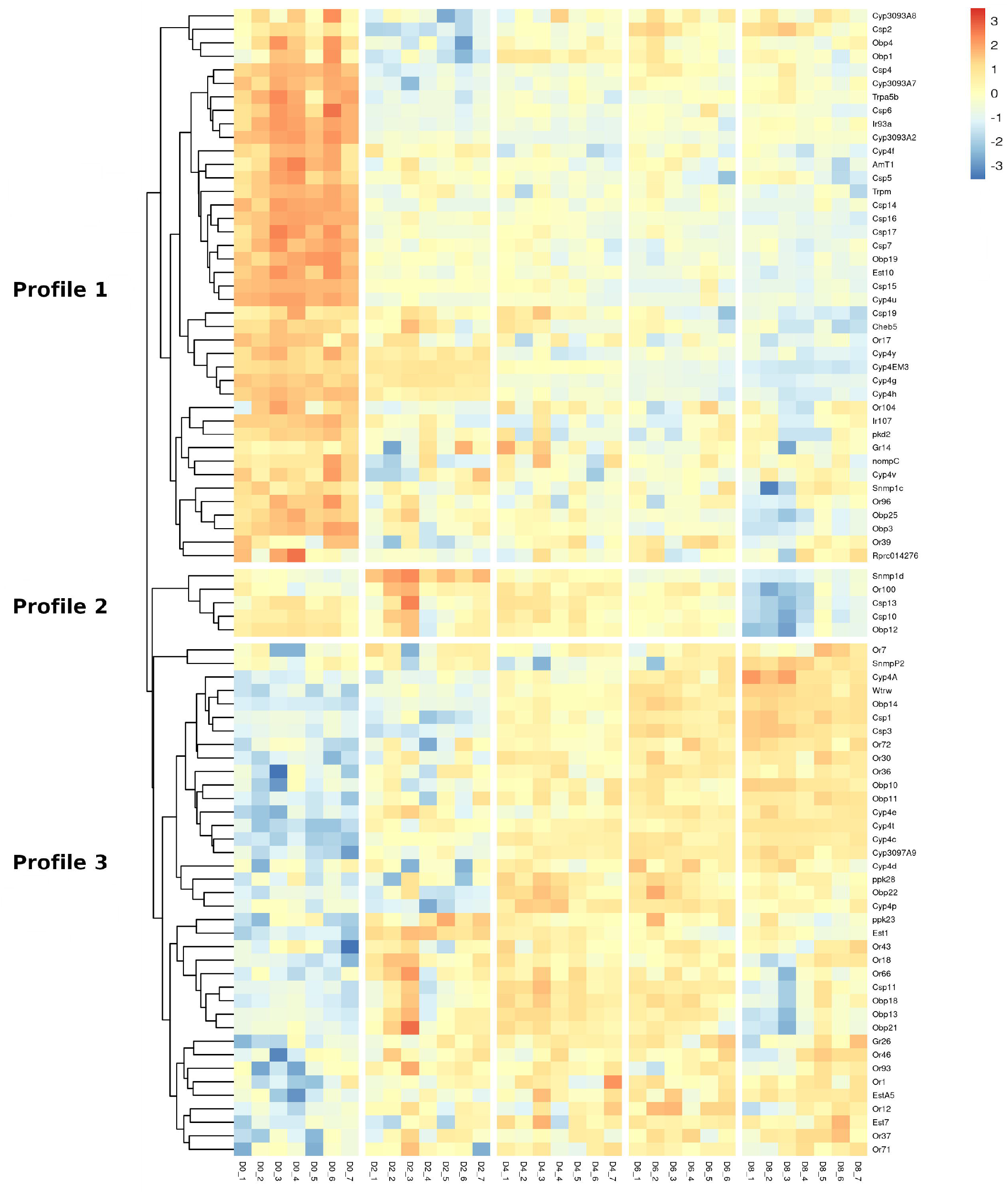
Box-plots of differentially expressed odorant binding and chemosensory proteins along five age points post-ecdysis. Gene expression is represented as Variance Stabilizing Transformed (VST) counts that were obtained from count data and the *vst* function from DESeq2. VST counts are reported in Additional file 10: Table S8.

### Expression changes of odorant degrading enzymes

A total of 4 secreted carboxylesterases esterases (Est) belonging to the pheromone/ hormone processing class and 17 members from cytochrome 4 (Cyp4) clade were classified as DEGs (Additional file 6: Table S4). Ten ODEs followed profile 1, showing very significant down-regulation with age, e.g., *RproCyp4EM3, RproCyp4g, RproEst10* or *RproCyp4h* (Fig. 6). In contrast, 11 ODEs fit into profile 3, showing a dramatically increased expression with age, e.g., *RproCyp4a, RproCyp4c, RproCyp4t, RproEst5* and *RproCyp3097A9* (Fig. 6).

**Fig. 6.**
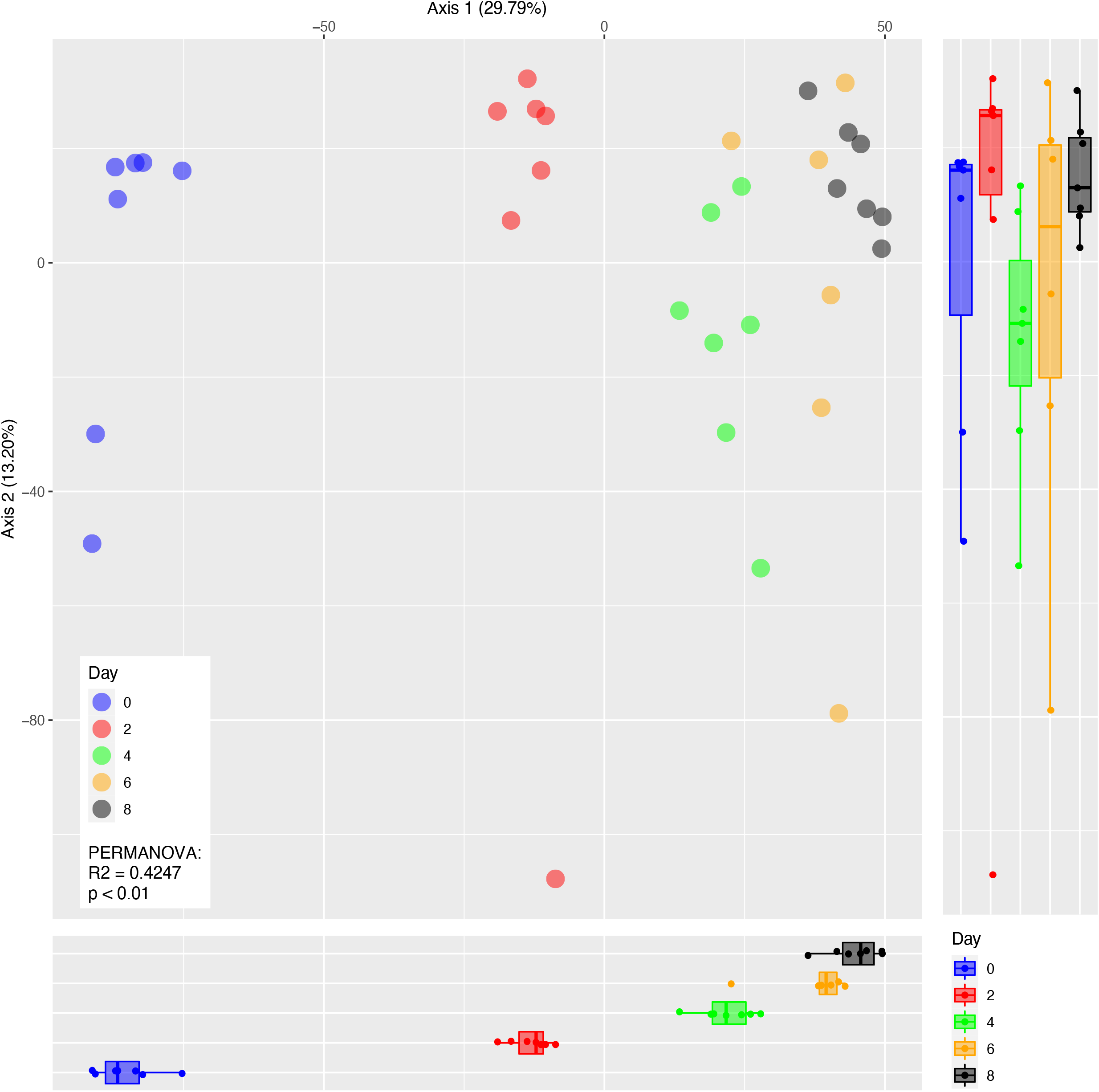
Box-plots of differentially expressed odorant degrading enzymes along five age points post-ecdysis. Gene expression is represented as Variance Stabilizing Transformed (VST) counts that were obtained from count data and the *vst* function from DESeq2. VST counts are reported in Additional file 10: Table S8.

## Discussion

Triatomine motivation for host searching is known to depend on age [30]. Nevertheless, it is not clear to which extent the central nervous system and peripheral sensory structures underlie the gradual increase seen through the first week after ecdysis. Our study shows that age induces dramatic changes in antennal gene expression that strongly correlate with the known age-dependent modulation of host search. Most of these transcriptional changes occur during the first two days after ecdysis, when more than 2,500 genes have their expression altered by more than 50%. The set of sensory genes having significantly increased expression along age may include potential candidates underlying the increased host cue responsiveness shown by *R. prolixus* eight days after ecdysis. This is the first time such a maturation process undergone by the peripheral olfactory system is described in an hemimetabolous insect.

The antennae of triatomines face major transcriptional expression changes during the first week, with a total of 6,120 genes (54,7 % of expressed genes) exhibiting significant modifications in their expression levels. The number of sensilla in triatomine antennae increases through development [31], supporting our results. In addition, part of these gene expression changes may relate to the known remodeling endured by sensilla during molting [32]. In all comparisons, a higher number of genes was significantly down-regulated, even though a relevant proportion of them underwent up-regulation (Table 1). This resembles the overall profiles described in the antennae of *Ae. aegypti* [8]. As shown by Fig. 2, there was no prevalence of downregulation in the case of differentially expressed sensory-related genes that present similar proportions of up/down-regulation. Indeed, the ratio between up and down-regulated genes varied between age point comparisons, e.g. between days 2 and 4 the genes that increased their expression were more frequent (Table 1).

The first two days after ecdysis seem to present the most dramatic modulation of antennal gene expression in *R. prolixus*. This is true both in terms of number of genes affected and intensity of expression changes (see PCoA and MA plots). We propose that most of this process is related to the structural transformation that antennae undergo during moulting transforming them at molecular and physiological levels. However, part of this change seems to be related to the remodeling of sensory function (Table 2). Interestingly, behavioral changes related to host-searching, e.g. the capacity to respond to CO2 or heat, appear later between days 6 and 8 [30]. It is difficult to propose candidate genes underlying this increased capacity to respond to host cues based on our results, as most sensory-related genes that showed significant modulation seem to change in the first two days after ecdysis. However, a reduced subset of genes increase their antennal expression through the first week, being candidates for functional genetics studies (Additional file 6: Table S4). Whether the peripheral sensory system is functional in molecular terms at the end of the age interval studied, contributing to host cues responsiveness, or if these behavioral changes are triggered centrally cannot be determined based on our results. Indeed, antennae seem almost identical at the transcript level between day 6 and 8 (only 80 DEGs, Table 1 and Fig. 1). A similar pattern has been observed for the antennae of *Ae. aegypti* [8] and *Anopheles coluzzi* [9], where age-dependent molecular changes occur before the behavioral switch.

As expected according to their functional role, olfactory co-receptors displayed stable expression (Additional file 7: Table S5). Consistently, *RproOrco* was the most expressed gene among all ORs, but *RproOr100* and *RproOr20* seem to deserve attention because they were the ORs showing highest expression. This is consistent with previous antennal transcriptomes from fifth instar nymphs and adults of *R. prolixus* [29]. Regarding IR co-receptors, *RproIr25a* was the most expressed followed by *RproIr8a* and *RproIr76b* (Additional file 7: Table S5), but transcripts of *RproIr93a* and *RproIr75a* were the most abundant also reinforcing the profiles reported by [29]. Interestingly, *Ir93a* has been described as a humidity sensor in *D. melanogaster* and a thermal receptor in *An. gambiae* [33], suggesting similar roles in triatomine host cue detection. The acetic acid responsiveness reported for *Ir75a* in *D. melanogaster* [34] suggests that detection of short-chain fatty acids may also be function of triatomine antennae. In fact, these compounds are known components of host odor, and in aggregation and alarm pheromones [15, 35, 36].

Triatomines and mosquitoes are not attracted by hosts right after ecdysis, but show consistent responses to them a few days later [9, 30]. Indeed, mosquitoes also present antennal expression changes in sensory genes that correlate with this behavioral plasticity [8, 9, 27] In our study, two profiles split most genes that show significantly modulated expression across age points (Fig. 2), suggesting opposing temporal regulation profiles for their sensory functions. Profile 3 genes are candidates to underlie the modulation of responsiveness to host-associated cues because their expression is directly correlated to the increase in behavioral responsiveness seen during the first days after molting [30]. We suggest that OSNs expressing the 13 ORs included in profile 3 (Fig. 3 and Additional file 6: Table S4) should present increased responsiveness to their cognate ligands at the end of the first week after ecdysis. Unlike ORs, the expression of GRs and IRs was very stable after ecdysis. Only one GR (*RproGr26*) presented a significant increase in expression with age, peaking at day 6 (Fig. 3). This gene together with two other phylogenetically-related GRs (*RproGr27* and *RproGr28*) are the most expressed genes of this family in bug antennae (Additional file 7: Table S5), suggesting that they possess fundamental roles for *R. prolixus* biology. We propose that the sensitivity of gustatory receptor neurons expressing *RproGr26* should increase in correlation with its transcript abundance.

Regarding OBPs, CSPs and ODEs, a total of 20 genes increased their expression with age (Fig. 5 and 6). The first two families of soluble proteins have been proposed to mediate the delivery of odor molecules to receptors found in OSN membranes [37]. On the other hand, it has been shown that ODEs degrade odor molecules accumulating in the sensillar lymph, allowing the clearance of already detected molecules [38]. Considering these functional roles, we suggest that the detection of compounds targeted by this set of proteins would become more efficient through the first week after ecdysis.

Only two PPKs (*RproPpk23* and *RproPpk28*) had their expression significantly increased as a function of post-ecdysis age (Fig. 4). In spite of the fact that a role in salt sensing has been described for these two PPKs in *D. melanogaster* [39] and *Ae. aegypti* [40], functional information is still scarce for triatomines. Nevertheless, it has been recently shown that *RproPpk28* detects high salt concentration in the context of host skin recognition [41], suggesting that this sensory ability may show increased sensitivity at the end of the first week post-ecdysis.

The TRP channel *water witch* (*wtrw*) is an hygroreceptor that detects moist air in *D. melanogaster* [42]. Host skin and the breath of vertebrate hosts are sources of water vapour that is used by triatomines to oriente at short-range during host seeking [43, 44]. It is possible that this function was conserved across insect evolution and that triatomines also use it as a hygroreceptor mediating host recognition. We suggest that its up-regulation across age points in *R. prolixus* antennae (Fig. 4) would support this idea and indicates that detection of water vapor is not fully functional right after ecdysis.

Sensory genes whose expression significantly decreased with age (profile 1) were mainly CSPs (10) OBPs (5) and ODEs (11). Four ORs (*RproOr17, RproOr39, RproOr96* and *RproOr104*), two IRs (*RproIr93a* and *RproIr107*), one GR (*RproGr14*), one PPK (*Rprc014276*) and four TRP channels (*RproNompC, RproPKD2, RproTrpA5b* and *RproTrpM*) presented this expression profile. Genes following this profile could mediate avoidance or behavioral inhibition to hosts as proposed for *Or39*, a receptor that detects human body odors that also presents a significant down-regulation between day 1 and 4 after molting in *An. coluzzi* [9]. An alternative hypothesis would be that the tasks related to these genes need to be fully functional right after ecdysis, having a relatively low turnover. Finally, it is known that the detection of alarm and aggregation signals is required at all ages and instars of triatomines. Genes that have high and stable expression in the antennae (Additional file 3: Table S3) might represent candidates for mediating similar functions.

## Conclusions

The antennae of *R. prolixus* undergo a great molecular remodelling during the first days after ecdysis, a process that very likely transforms them at the physiological level. Part of this transformation is reflected in the transcript abundance of several sensory receptors and olfaction-related genes. We were able to identify a set of genes with sensory function whose expression increases across age, therefore becoming candidates to mediate the detection of host cues in this Chagas disease vector. However functional genetics studies, using RNA interference, CRISPR-cas9 technology or heterologous expression systems to allow their “deorphanization” are still necessary to transform them into feasible targets to manipulate triatomine behavior in the future.

## Methods

### Insects and RNA isolation

*Rhodnius prolixus* were reared under controlled conditions (26±1°C temperature, 65±10% relative humidity and a 12h:12h light/dark cycle) at the Instituto René Rachou - FIOCRUZ. Antennal samples were obtained from unfed fifth instar larvae along five different age points (0, 2, 4, 6 and 8 days) after ecdysis. Each time point was replicated 7 times using 20 antennae per sample. Antennae were manually homogenized using sterilized pestles and total RNA was extracted using TRIzol® Reagent (Life Technologies, Carlsbad, CA, USA) according to the manufacturer’s instructions. Then, extracted RNA was resuspended in 22 μL of DEPC-treated water (Life Technologies), and its concentration determined at using Qubit (Thermo-Fisher Scientific). RNA integrity and quality were assessed by means of a 1% agarose electrophoresis gel and in Agilent 2100 Bioanalyzer (Agilent Technologies, Santa Clara, CA, USA).

### Illumina sequencing an quality control

Library construction and sequencing services were performed by the Max Planck Genome Center (Cologne, Germany). RNA-Seq libraries were constructed using the TruSeq mRNA Sample Preparation Kit (Illumina, San Diego, CA). A total of 35 libraries were sequenced using Illumina HiSeq2500 from both ends of the molecules to a total read length of 150 bp from each end. The raw sequence dataset is available with the NCBI-SRA Bioproject number PRJEB44111. The sequencing output has higher than 15 million reads in all libraries (Additional file 8: Table S6). The FASTQC software (bioinformatics.babraham.ac.uk/projects/fastqc/) was used to assess the read quality and the presence of Illumina sequencing adapters. Afterward, Illumina adapters and those bases from 5’ and 3’ ends with quality scores lower than 5 (TRAILING: 5 and LEADING: 5 parameters) were eliminated from the reads using Trimmomatic v0.32 in the paired-end mode [45]. The SLIDING-WINDOW parameter was fixed at 4:15 and only those reads longer than 50 base paired were kept for the next steps.

### Mapping reads and differential gene expression analysis

The trimmed reads (between 14 and 12 million reads per library) were mapped to the *R. prolixus* genome assembly (version RproC3.3 from VectorBase) by means of STAR v.2.6.0 [46] with default parameters and an edited General Feature Format file generated by [47]. The percentage of mapped reads was between 71 and 79% in all libraries, except for one replicate from day six with only 11% of mapped reads. Subsequently, the *multicov* command from the BEDTools suite v.2.27.0 was used to report the count of alignments per feature (Additional file 9: Table S7).

### Highly expressed genes along age

The expression of each target gene was estimated as Transcripts Per Million (TPM) for each replicate and the geometric mean for each time point was calculated (Additional file 7: Table S5). Expression values of target genes were ranked for each time point and the top 50 genes were extracted. Finally, a consensus list of target genes present in the five rankings was elaborated.

### Differentially gene expression analysis

Global analysis – EdgeR v3.30.1 package [48] was used to detect global changes across time in *R. prolixus* antennae. After the normalization (using *calcNormFactors* function) and estimation of gene-specific biological variation (using *estimateDisp* function) steps, normalized counts were analyzed using the GLM approach and the quasi-likelihood F-test. DEGs were identified considering a total of eight comparisons: sequential (day 0 *vs*. day2, day 2 *vs*. day 4, day 4 *vs*. day6 and day 6 *vs*. day 8) and grouped (day 0 *vs*. all, days 0-2 *vs*. days 4-6-8, day 0-2-4 *vs*. day 6-8, all *vs*. day 8). Genes with low expression and/or high variation were filtered using the HTSFilter v.1.32 [49]. Those genes with an FDR<0.05 were considered as differentially expressed.

Pairwise comparisons - DESeq2 v.1.28.0 [50] was used to perform pairwise comparisons between the time points and groups mentioned above. First, the HTSFilter package was used to filter gene counts for each comparison. Following, each filtered dataset was used to identify DEGs using the *apeglm* estimation from the *lfcShrink* function. This function is based on the “Approximate Posterior Estimation for Generalized Linear Model’’ (*apeglm*) method that uses an adaptive Bayesian shrinkage estimator to generate more accurate log2 fold-change values [51]. Finally, a DEG list was generated for each comparison based on *s*-values <0.05 to define the significance level and on a fold-change threshold >1.5. The output generated by the *lfcShrink* function was also used to elaborate MA-plots that show the mean of the normalized counts versus the log2 fold-changes for all genes tested.

Count data were transformed using the Variance Stabilizing Transformation (VST) for negative binomial data with a dispersion-mean trend [52], implemented in the *vst* function of DESeq2 (Additional file 10: Table S8). Following, transformed values of DEGs were extracted and used to generate heatmaps (using the pheatmap package in R) and box-plots. The transformed values of transcript expression were also used to calculate the euclidean distance between libraries and create a Principal Coordinate Analysis (PCoA) plot using the *ade4* R package [10.18637/jss.v022.i04]. The impact of the variable day on the distances was measured by a PERMANOVA analysis using 999 permutations in R.

## List of abbreviations

SNs: Sensory neurons
OSNs: Olfactory sensory neurons
GLM: General Linear Model
DEGs: Differentially expressed genes
OBPs: Odorant binding proteins
CSPs: Chemosensory proteins
ODEs: Odorant degrading enzymes
ORs: Odorant receptors
SNMPs: Sensory neuron membrane proteins
IRs: Ionotropic receptors
GRs: Gustatory receptors
PPK: pickpockets
TRP: Transient receptor potential
Est: Secreted esterases
Cyp: Cytochromes
GRNs: Gustatory receptor neurons
TPM: Transcripts Per Million
VST: Variance Stabilizing Transformation
PcoA: Principal Coordinates Analysis

## Declarations

### Ethics approval and consent to participate

Not applicable

### Consent for publication

Not applicable

### Availability of data and materials

All data generated or analysed during this study are included in this published article and its supplementary information files.

### Authors’ contributions

M.G.L conceived the project. M.G.L, E.G.W, G.R.F, B.S.H. and J.M.L.E designed the experiments and performed data analyses. J.M.L.E. generated insects and RNA samples for RNA-Seq. J.M.L.E, E.G.W and G.R.F carried out the bioinformatic analyses and provided RNA-Seq data. All authors wrote the manuscript and provided comments on versions. All authors read and approved the final manuscript.

## Acknowledgements

The authors acknowledge the Bioinformatics Platform of René Rachou Insitute for providing the computational resources to perform the analyses.

## Additional files

**Additional file 1: Table S1. Results of PERMANOVA Pairwise comparisons.** Df: Degrees of freedom;

**Additional file 2: Table S2. Results of the differential gene expression analysis using the GLM approach performed in EdgeR.** Fc: Fold change; CPM: Count per million; FDR: False Discovery Rate.

**Additional file 3: Table S3. Sensory genes showing highest expression along 8 days post-ecdysis.** This list was generated from the consensus of the 50 most expressed sensory genes in each time point as described in Supplementary Table S5. Gene expression (expressed as TPM) was calculated using the geometric mean of the five time points (obtained with the geometric mean of each technical replicate *per* time point). TPM: Transcripts Per Million; DEG: Differentially Expressed Gene.

**Additional file 4: Fig. S1. MA-plots of Day 0 *vs*. Day 2 (top left), Day 2 *vs* Day 4 (top right), Day 4 *vs* Day 6 (bottom left) and Day 6 *vs* Day 8 (bottom right) representing log fold-change versus mean expression of normalized counts.** Differentially expressed genes are shown in red. The blue line represents a logFold change cut-off (0.583, equivalent to a 50% fold-change) used to identify differentially expressed genes (DEGs).

**Additional file 5: Fig. S2. MA-plots of Day 0 *vs*. All (top left), Days 0 and 2 *vs* All (top right), Days 0,4,6 *vs* Days 6 and 8 (bottom left) and Day All *vs* Day 8 (bottom right) representing log fold-change versus mean expression of normalized counts.** Differentially expressed genes are shown in red. The blue line represents a logFold change cut-off (0.583, equivalent to a 50% fold-change) used to identify differentially expressed genes (DEGs).

**Additional file 6: Table S4. Results of the differential gene expression analysis performed in DESeq2.**

**Additional file 7: Table S5. Transcripts Per Million values for sensory genes in the 34 libraries and geometric mean for each time point (green) and the five time points.**

**Additional file 8: Table S6. Raw, trimmed and mapped reads, including percentages, for each library.**

**Additional file 9: Table S7. Raw counts for each gene in the 35 libraries.**

**Additional file 10: Table S8. Variance Stabilizing Transformed counts for each gene in the 34 libraries.**

