## Supplementary figures and images for "Changes in antennal gene expression underlying sensory system maturation in *Rhodnius prolixus*"

### Supplementary Figure 1

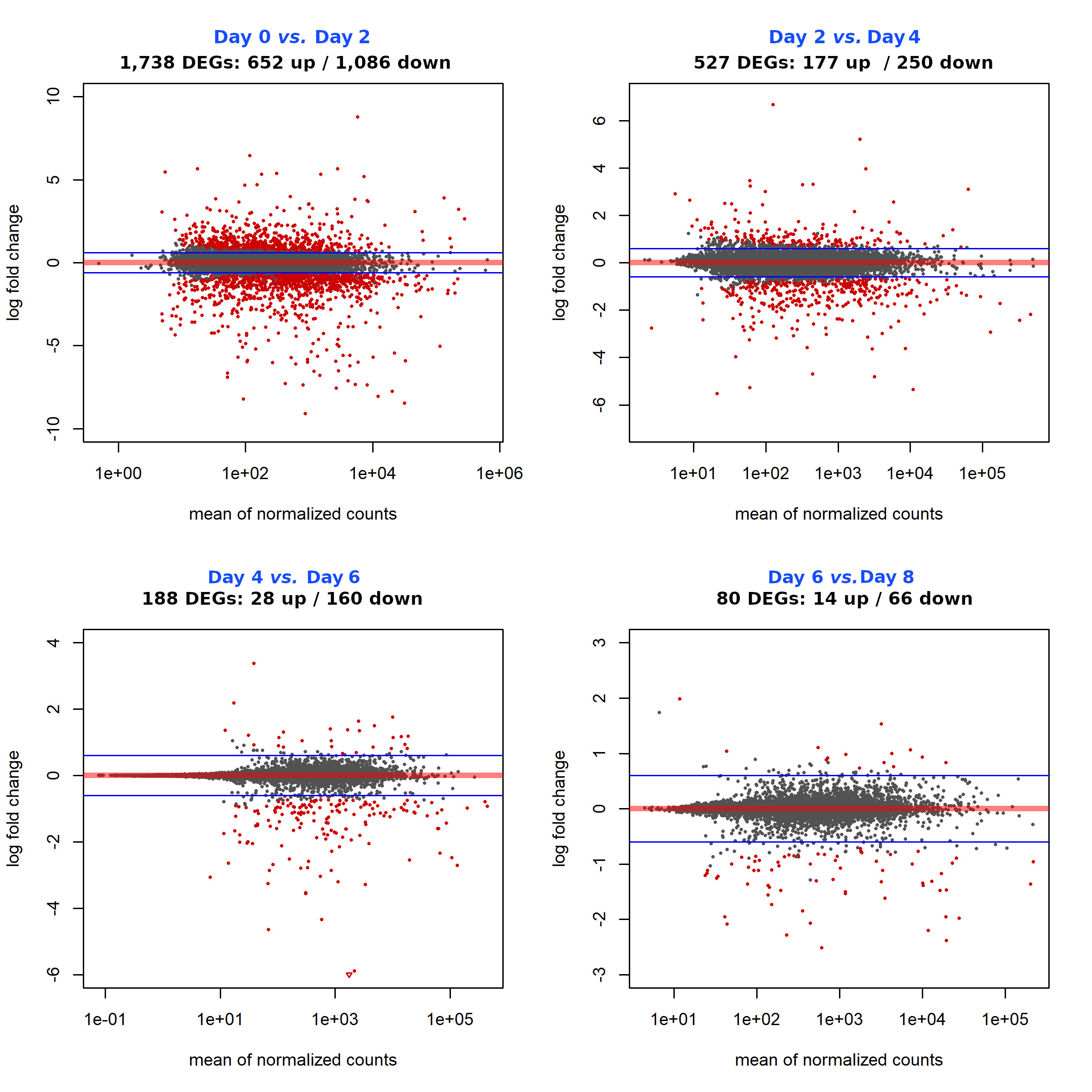

### Supplementary Figure 2

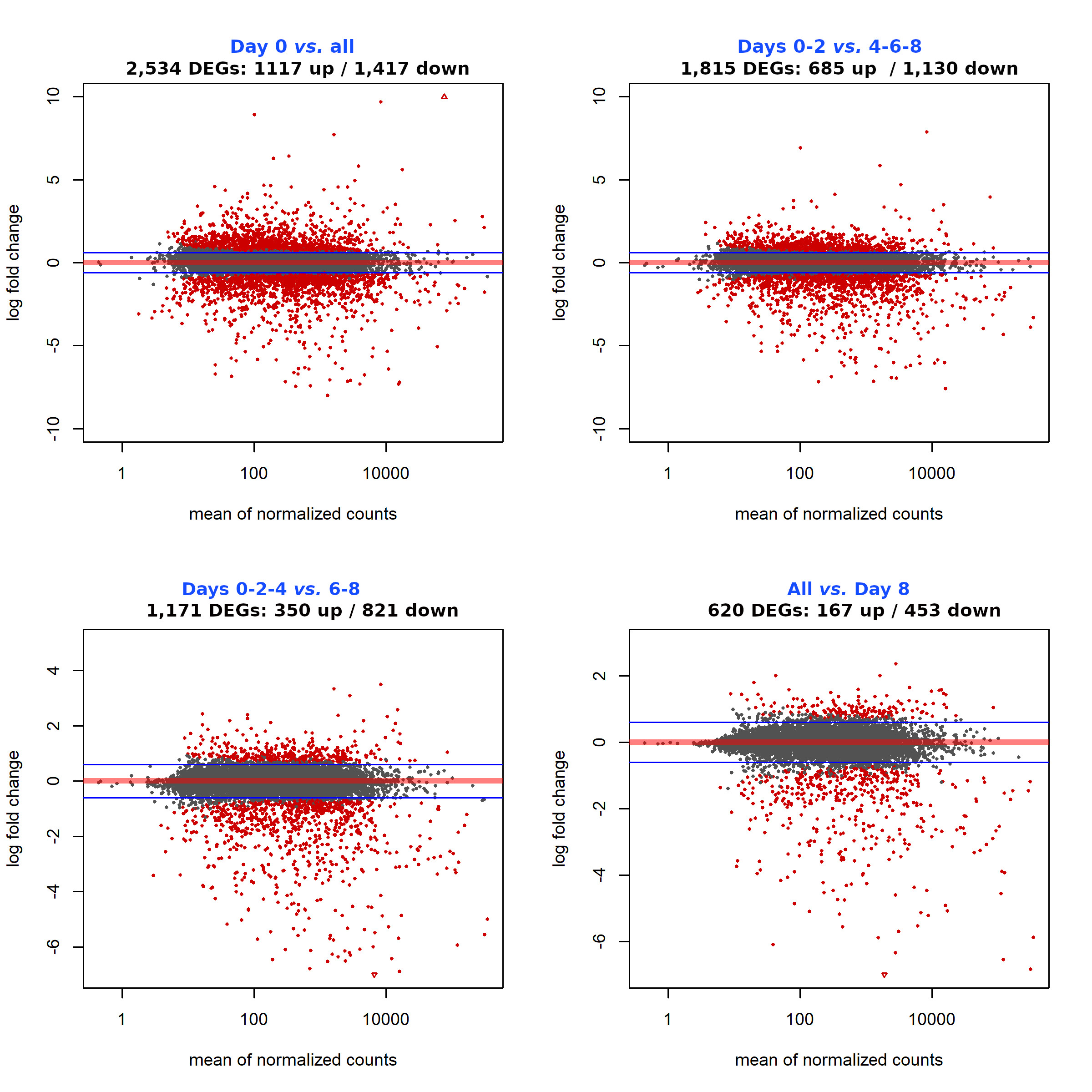
